# A mechanistically-inspired geometric model to predict microbial growth across environments

**DOI:** 10.1101/2025.07.02.662826

**Authors:** Thomas Tunstall, Urszula Łapińska, Stefano Pagliara, Krasimira Tsaneva-Atanasova

**Affiliations:** Living Systems Institute, University of Exeter, Stocker Road, Exeter, EX4 4QD, Devon, UK; EPSRC Hub for Quantitative Modelling in Healthcare, University of Exeter, Stocker Road, Exeter, EX4 4QJ, Devon, UK; Biosciences,University of Exeter, Stocker Road, Exeter, EX4 4QD, Devon, UK

**Keywords:** Theoretical Ecology, Bacterial Growth, Mean-field Model, Population Dynamics

## Abstract

Estimating bacterial growth trajectories is essential for bridging mechanistic insights and experimental data. Nevertheless, widely used empirical models tend to be overly simplistic or purely phenomenological. Mechanistic models offer greater detail but still fall short of representing the full spectrum of biochemical processes underlying cellular growth. Here we take a mean-field approach whereby we consider the bacterial cell population as a cascading sequence of biochemical processes. This approach enables the derivation of a geometry-based model of cellular population growth that captures the lag, exponential, and stationary phases observed experimentally, without requiring assumptions about specific cellular mechanisms. Furthermore, parameters estimated from multiple datasets can provide insights into the population’s history of cellular stress. We demonstrate the robustness and accuracy of the proposed modelling framework by applying it to existing experimental data for *S. aureus* and *P. aeruginosa* grown in closed and open environments. We show that our approach achieves higher accuracy compared to contemporary models.

## 1 Introduction

Bacterial growth in the presence of limiting resources is characterised by four main phases [1–3]: the lag phase, in which there is little growth of the bacterial population, the exponential phase, in which the population increases at an exponential rate, the stationary phase, where the population plateaus, and the death phase, where the population decays. In a closed system, from which bacteria cannot escape and nutrients cannot be resupplied, plotting population density over time produces a characteristic sigmoidal curve that reflects the first three growth phases. This curve is most commonly fitted using logistic or Gompertz functions [4–7]. Such fitting is phenomenological as opposed to mechanistically derived, and is usually based only on symmetry properties of the observed data. The logistic function is employed when the growth curve is symmetric, and the Gompertz function is often used when there is asymmetry between the transition from the lag to the exponential phase and the transition from the exponential to the stationary phase.

Mechanistic models of bacterial growth necessitate a complete or partial understanding of all the relevant metabolic pathways and biochemical processes within each cell in a bacterial population, which can vary between species, strains as well as different cells within the same bacterial population [8]. Such complexity is exponentiated when we transition from growth in monoculture to growth in coculture: antagonistic and synergistic interactions increase the number of biological pathways that need to be considered [9]. It is therefore useful, for both fitting and data comparison, to derive a minimal model based on mechanistic underpinning. Such a model ought to capture the main phases of bacterial growth, accounting for the history of stressors experienced by the cells.

As a modelling assumption, we take the view that each cell consists of a high number of high-level biochemical processes - or ‘components’ - the products of which are used in lower order biochemical processes resulting in biomass production. Thus, if a single component is rendered inactive, for example because of nutrient unavailability, the presence of toxic compounds, or cellular transition to a dormant state, all downstream processes dependant upon those products will be rendered inactive. In this way, we have employed a mean-field approximation of the complex cascade of intracellular processes to derive a geometrically-based, effective cellular replication rate which yields a general, minimal, mechanistic model of bacterial population growth.

To demonstrate the applicability of the model, we apply it to recently described data (see Ref [10]) of *S. aureus* and *P. aeruginosa* grown in monoculture and coculture, in both closed systems (where nutrients can become limiting) and open systems (where nutrient availability is non limiting). We further compare monoculture growth in closed systems with commonly used fitting functions and a kinetics-based model.

## 2 Model

Even the simplest biological cell is a complex dynamical system, consisting of many biochemical processes working in concert to facilitate cellular replication [11]. To simplify nomenclature, we shall refer to relevant biochemical processes and specific organelles or molecules as ‘components’. Components can be rendered inactive, either passively by resource unavailability, or actively due to inhibition from the cell itself. For example: when a resource is limiting, components within a cell are passively rendered inactive as necessary substrates are unavailable. Cells may also actively enter a state of dormancy to survive periods of resource scarcity [12] or toxicity build-up (such as persister cells in the context of resistance to antimicrobials [13–15]): in these cases, the cell may be effectively rendering its own components inactive. We shall construct a mean-field model by mapping the consequence of component inactivity for an idealised average cell, i.e. that does not account for cellular heterogeneity [16], which maps to the full population.

Fig. 1a illustrates the consequences of a component being rendered inactive during an infinitesimal time-frame: if all components are active, all downstream processes are able to take place. On the other hand, if a component is not active then all downstream processes depending on the products of that component are rendered inactive too. We assume that the rate of cellular growth is proportional to the fraction of active downstream processes. To gain a better geometrical understanding, we construct a hypothetical ‘interaction-space’, consisting of all biochemical reactions which happen within a small period of time within an optimally functioning cell (illustrated in 2D in Fig. 1, right). If a single component is removed (component 1), this causes the shut down of all downstream reactions dependent upon it, effectively carving out a volume *β* from interaction-space. The effective growth rate of the average cell is the proportion of interaction space which is active. If another component is removed (either component 2 or 3), there are two possibilities: the removal of component 3 can have consequences independent of the first, as an independent suite of downstream processes are shut down. On the other hand, if component 2 is rendered inactive the impact is mitigated as it shares downstream processes which have already been disabled by the inactivation of component 1. Geometrically, this is akin to carving out another volume of interaction-space, which may overlap with the first. Given the remaining fraction of interaction space corresponds to the modulation of the growth rate, and that we consider the limit of many components, we can use known geometrical and statistical results [17, 18] to assert that, given a proportion of inactive components *I*, the growth rate of cells is *r*(*t*):

**Fig. 1:**
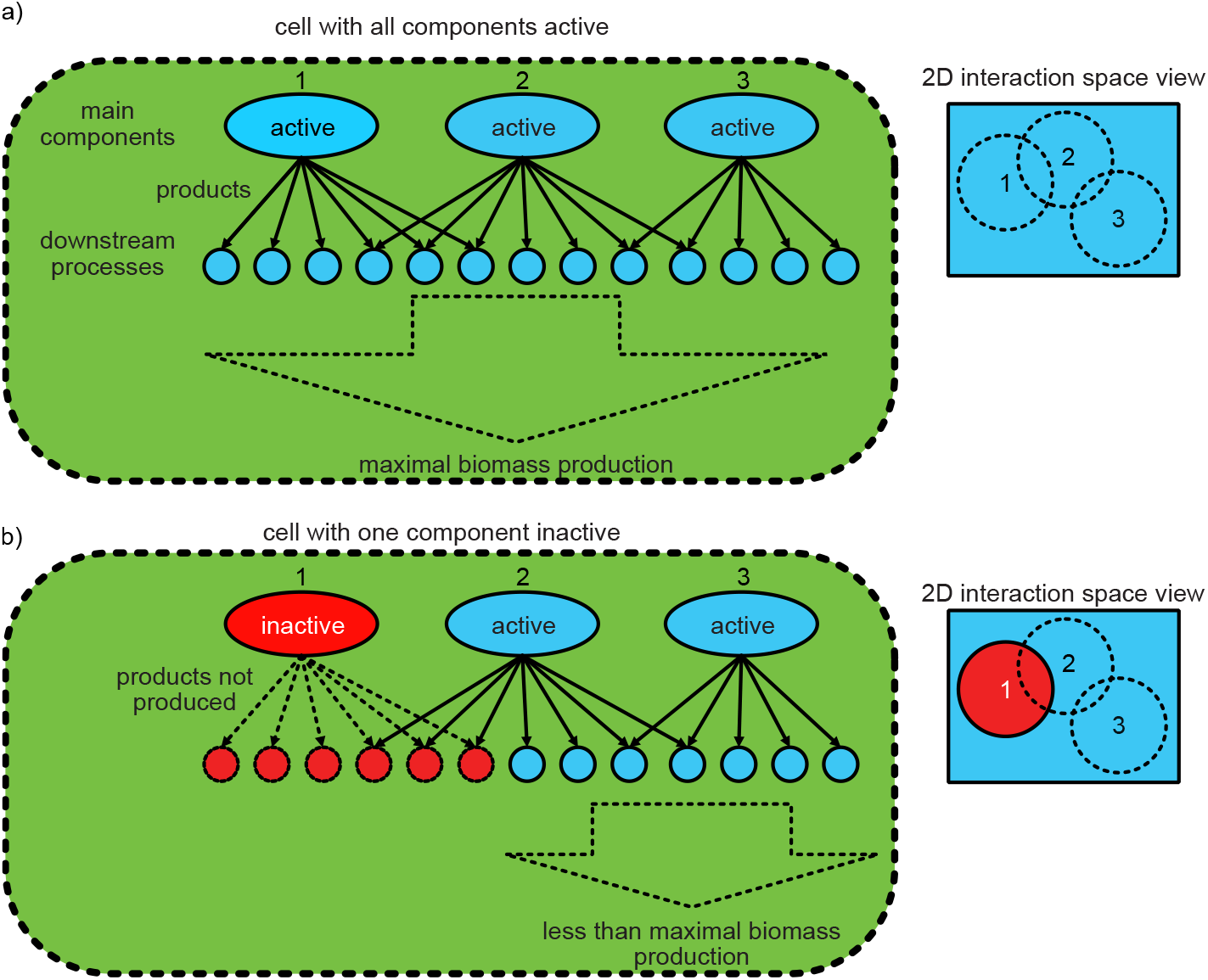
Illustration of the consequences of intracellular components being rendered inactive. Only three components are considered, and the products of these components are used in downstream processes which result in biomass production. **a)** All high-level components are active, meaning products of these components are able to be utilised in downstream processes to generate biomass at an optimal rate (left panel). In the 2D interaction-space perspective (right panel), the components 1, 2 and 3 correspond to circles of radius *β*. **b)** When a single component (e.g. component 1) is rendered inactive, the downstream processes dependent upon the products of this component are also rendered inactive, resulting in a sub-optimal rate of biomass production (left panel). In the 2D interaction-space perspective (right panel), we see that the removal of component 1 carved out a volume *β* from interaction-space. Note that the subsequent removal of component 3 would have an independent impact on the growth rate as there are no shared downstream processes. However, if component 2 were removed, the consequence is less than additive as 1 and 2 share downstream processes in interaction-space.

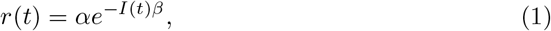

where *α* is the maximum growth rate, and *β* is a measure of the downstream consequence of component inactivation. Note that in this formulation, as *I* tends to 1, the growth rate tends not to 0 as we would physically expect, but to *e*^*-β*^. This is a consequence of the assumption of spatio-temporal correlation in interaction space: we have implicitly assumed that nearby components share nearby downstream processes. We can apply a shift and re-scale (an affine transformation) to ensure this biologically-realistic condition is met:

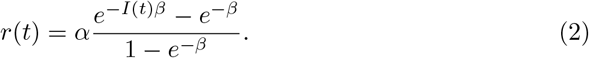

This formulation can also be derived probabilistically by assuming that the products contribute independently to downstream processes which is analogous to assuming that the products are well-mixed within the cell. (see Appendix A).

Next, we shall consider how the fraction of inactive components within the average cell varies over time: the motivating example shall be bacterial growth in a closed environment, first emerging from the lag phase.

When a cell that has become inactive because of nutrient unavailability restarts receiving nutrients, the cell begins activating inactive components independently from each other. We shall assume that there is one variety of component which has the most dramatic impact on downstream biomass production: the net rate of activation of these components shall be *γ*_1_. Assuming that the initial fraction of inactive components is *I*_0_, the portion of inactive components thus initially evolves over time according to:

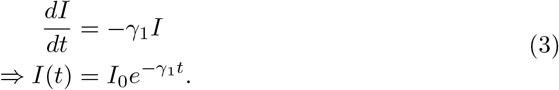

We highlight that *I*_0_ is effectively a parameter that accounts for the history of stress experienced by the cell [19], and thus is difficult to estimate *a priori*.

If nutrients do not become limiting, we note that lim_*t→∞*_*I*(*t*)*→*0, and thus *r*(*t*)*→α* and the population remains in the exponential phase. However, if the population grows in a closed environment, some nutrients will eventually become limited, and/or a waste product will increase the probability that cells become inactive. Here, we assume for simplicity that nutrients becoming limiting is the main issue. In a population undergoing exponential growth, nutrients become limiting in a finite time [20]. Therefore, as a simplifying assumption, we also assume that a nutrient becomes a limiting factor at time *T*. *T* is therefore a parameter to account for the rate of nutrient uptake, as well as the initial concentration of that nutrient. Given a vital nutrient is no longer present, e.g. glucose [20], components become inactive at a constant rate *γ*_2_, either as a result of components running out of the essential nutrient, or by the cell beginning to enter a state of inactivity. Thus, the overall growth of the population obeys the following equation:

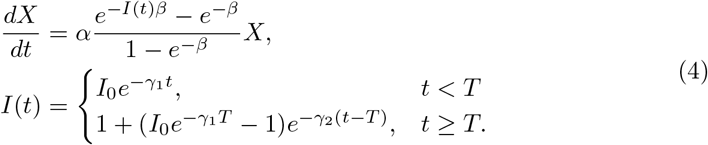

### 2.1 Model Analysis And Interpretation

Our mean-field model can be used to discern particular transitions between stages of cellular growth at the population level: from the lag phase to the exponential phase, and from the exponential phase to the stationary phase. Furthermore, we can readily make adjustments to account for the differences between a closed environment and an open environment.

#### 2.1.1 Closed Environment

The term closed environment refers to standard bacterial culturing in well-mixed flasks, where the bacterial inoculum and nutrients are added at the onset of the culture and are not resupplied in time: in these systems bacterial growth leads nutrients becoming limiting in finite time [20].

Although the lag phase has been long thought to allow cells to adapt to new environmental conditions by adjusting their metabolism and gene expression, thereby enabling subsequent growth [3], recent advanced proteomics analysis [21, 22] demonstrate that during this phase the cells alter their physiology to enter reproduction: a comprehensive review of mechanistic models which generate a lag phase is found in Ref. [23]. Of particular interest are the models which implicitly account for the growth rate of bacterial cells by modelling how essential substances (such as enzymes, signalling molecules, etc.) are produced and metabolised within the cell. The Baranyi model [23–25] recovers a distinct lag phase by modelling the development of essential intracellular regulators within the cell with Michaelis-Menten kinetics, resulting in a single algebraic expression. The synthetic chemostat model (SCM) proposed in Ref. [26] recovers a distinct lag phase by employing a system of four ordinary differential equations to account for the concentrations of substrates, monomeric intermediates, and biomass within the cell. In essence, whereas the transition to the stationary phase is determined by environmental limitations, the lag phase is a property of the bacterial strain and the history of stress experienced: in this way, the transition from the lag phase to the exponential phase is a more fundamental property of the average cell compared to the transition from exponential to stationary phase. In light of this, the subsequent discussion will predominantly centre on the lag phase.

To discuss the predictions of the model regarding the lag phase and exponential phase, we shall consider only the time where *t < T* : this corresponds to the bacteria growing in an environment in which no resources are limiting. The effective equation is thus given by:

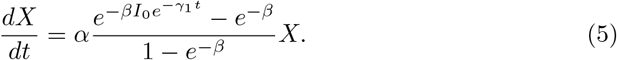

Here, we demonstrate that the growth rate is modulated according to an affine transformation of a Gompertz function. Historically, the Gompertz function has been used as the growth curve when there is asymmetry between the transition from lag to exponential phase and from exponential to stationary phase [7]. The Gompertz equation falls out from the Gompertz law [27], which describes the rate of fatality in the actuarial context. As it has been previously remarked in Ref. [28], the Gompertz law arises in many complex, inter-dependent systems: the key difference here is that instead of system failure causing death, it causes a failure to replicate in bacteria. Thus, we have shifted the Gompertz equation from a growth curve to a growth rate.

Eq. (5) can be solved:

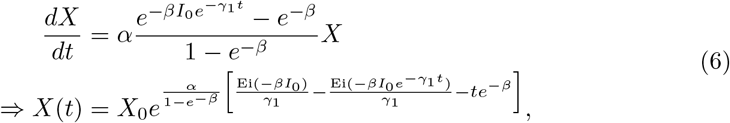

where Ei(*x*) is the exponential integral function. We note that Eqn. 6 exhibits the behaviours expected of a bacterial population transitioning from lag to exponential phase. When *γ*_1_*t* is small, the result approaches a constant value of *X*_0_. When *γ*_1_*t* is large, we recover exponential increase modulated by a constant factor.

The duration of the lag phase, *t*_lag_, can be characterised by the maximum of the second derivative of the natural logarithm of the growth curve [6], i.e.:

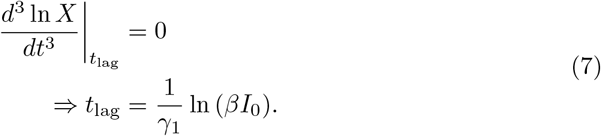

We note that *t*_lag_ also corresponds to the time of maximum slope in the Gompertz equation which acts as the modulation to the growth rate in Eq. (5). We can estimate the ‘width’ of the transition from lag to exponential phase by similarly solving for the maximum of the third derivative of the log of the growth curve: these maxima occur for 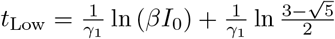 and 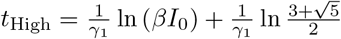. The width of this transition, *w*, is thus 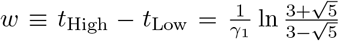 : naked-eye observations of the time at which the lag phase ends, and the transition time between phases, thus enable estimates of *βI*_0_ and *γ*_1_.

The analytical forms of *t*_lag_ and *w* allow us to map between the mechanistic and practical understandings of *γ*_1_ and *β*: mechanistically, *γ*_1_ corresponds to the rate of component reactivation, and this translates to controlling the duration of the transition from lag to exponential phases. Mechanistically, *β* corresponds to the impact of inactive components on the proportion of active biochemical processes in the cell, and this translates to controlling the location of the end of the lag phase.

Next, let us consider the transition of the population from exponential to stationary phase: we assume that the proportion of inactive components within the mean-field cell approaches zero prior to the onset of nutrient limitation. In this case, we consider:

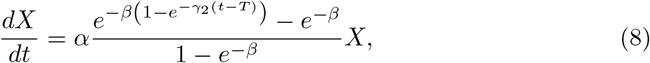

which can be solved similarly to Eq. (5) to derive characteristic timescales corresponding to the end of the growth phase. The transition from maximum growth to slow growth is commonly captured by modulating the maximum growth rate by the Monod function [3], which was first described phenomenologically as a function of the concentration of external substrate, *S*_*e*_, but it has since been derived mechanistically [29, 30].

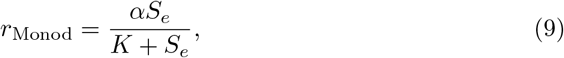

where *K* is the concentration of substrate at which the growth rate is half the maximum rate. The geometric growth rate is able to capture both the lag and stationary phases alone, while the Monod rate is only capable of capturing the stationary phase, and the fact that the geometric model is exactly solvable makes it computationally quicker and easier to evaluate.

#### 2.1.2 Open Environment

With open environment we refer to bacterial culture carried out in systems such as chemostats [31] or microfluidic devices (such as in Refs. [8, 14]) where bacteria are added to the systems at the onset of the experiments and bacterial growth is modulated by controlling the environment, such as by removing excess bacteria and bacterial waste while supplying nutrients. In these systems it is possible to avoid bacteria entering the stationary phase, and thus removing the need for *T* and *γ*_2_ to be accounted for: in the absence of other considerations, Eq. (6) describes the evolution of populations.

In this type of systems, motile organisms, such as *P. aeruginosa* are capable of escaping the system. The greater the motility of the organisms, the higher the rate of escape. Motility of *P. aeruginosa* is controlled by the *las* and *rhl* quorum-sensing systems [32]: these systems are affected by a variety of molecules, from intracellular molecules, such as cyclic di-GMP (lower concentrations corresponding to greater motility) [33, 34] to extracellular molecules such, as nitric oxide [35] produced due to oxidative stress (higher concentrations correspond to higher motility) and nutrient availability [32]. As a simplifying modelling assumption, we shall assume the quorum-sensing of *P. aeruginosa* is predominantly mediated by a single chemical (of concentration *Q*(*t*)) secreted by *P. aeruginosa*, and that the concentration of this chemical stimulates *P. aeruginosa* to increase motility, and thus escape the open system faster. From experimentation, in Fig. 2, we see that a minority of *P. aeruginosa* cells leave the microfluidic chambers when cells are loaded in the chambers from an overnight stationary phase culture. This observation suggests that the secretion of the quorum-sensing chemical and/or the motility of the cell is dependent upon the overall activity of the cell: as another minimal working assumption, we shall describe these in the same way that cellular growth is. The simplest set of equations describing this system is the following:

**Fig. 2:**
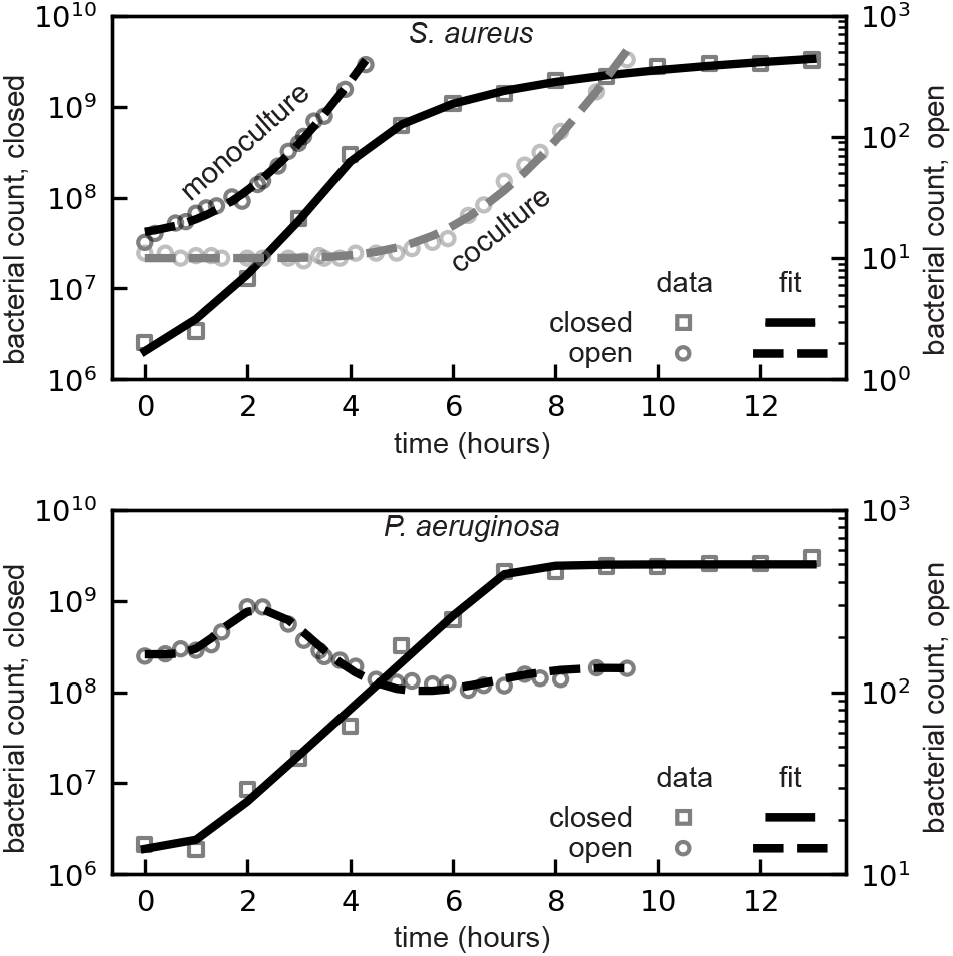
Comparison between data from Ref. [10] for open (circles) and closed systems (squares) to Eq. (4) for *S. aureus* (top), and Eq. (4) and Eq. (10) for *P. aeruginosa* (bottom). Fitting for closed system is a solid line, and open systems are dashed. Parameters are in Table. 1.

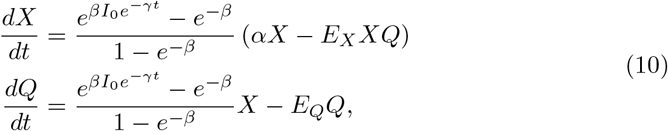

where *E*_*X*_ is the rate constant of *P. aeruginosa* escape, and *E*_*Q*_ is the rate constant of the quorum-sensing chemical escape. From here, we can analyse the steady state of the system. In the long term, the exponential factors tend to 1:

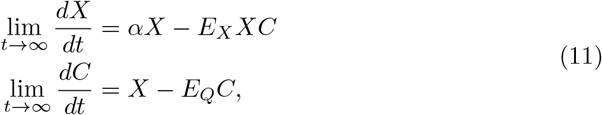

the stable point of which, (*X*^*∗*^, *L*^*∗*^), satisfies:

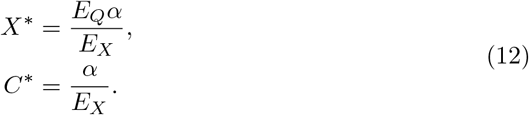

The eigenvalues about this point are 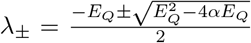 : given *E*_*Q*_ *>* 0, this is a stable steady state with damped oscillations for 4*α > E*_*Q*_, and a non-oscillatory steady state otherwise. The period of the oscillatory state is 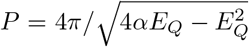; given the maximum growth rate, *α*, of *P. aeruginosa* is known from experimentation, the parameter *E*_*Q*_ can be inferred by measuring the period of oscillation.

## 3 Model Evaluation And Comparisons

To demonstrate the model’s capacity to describe bacterial growth, we fit it to the growth data of two bacterial species, *S. aureus* and *P. aeruginosa*, grown in mono-culture in both closed, i.e. well-mixed flasks, and open environments, i.e. microfluidic devices [10]. Due to the finite resources in a closed system, growth ultimately leads to nutrient limitation, and thus we expect that bacteria enter the stationary phase of growth. Hence Eq. (4) represents the appropriate model to employ for both species grown in monoculture. In the open system, where no nutrient becomes limiting for the duration of the experiments, Eq. (5) is appropriate for the non-motile *S. aureus* and Eq. (10) for the motile *P. aeruginosa*.

We assume that the relevant components rendered inactive in the cases of *S. aureus* in monoculture in closed systems are a result of nutrient unavailability: we therefore assume that *α, β*, and *γ*_1_ are the same in the open and closed cases, with only the initial conditions (the initial cell concentration, *X*_0_, and the initial proportion of inactive components, *I*_0_) differing between experiments. As *P. aeruginosa* is known to have an antagonistic effect on *S. aureus* [9, 10, 36, 37], we expect that the relevant components endered inactive may be different compared to those rendered inactive due to nutrient starvation, thus *β* and *γ*_1_ ought to differ for the co-culture open system of *S. aureus*. A further consequence of this interaction is that there is no simple data-driven method for discerning the *I*_0_ and *β* parameters, making it difficult to obtain an initial guess for least-squares fitting: to resolve this, in the *S. aureus* open co-culture, we employ the non-corrected growth rate Eq. (1) (recall that this model yield identical quantitative features for the lag-exponential phase transition). On the other hand, as there are no known relevant antagonistic effects on *P. aeruginosa* from *S. aureus* we assume that *α, β*, and *γ*_1_ are the same for *P. aeruginosa* in open and closed systems, in both monoculture and co-culture.

Fitting of the mechanistic models using least-squares regression (using `scipy.optimize.least_squares` [38]) requires an initial estimation of the parameters: in order to obtain the fitted parameters in Table 1, we employ the estimations discussed in Appendix B.

**Table 1:**
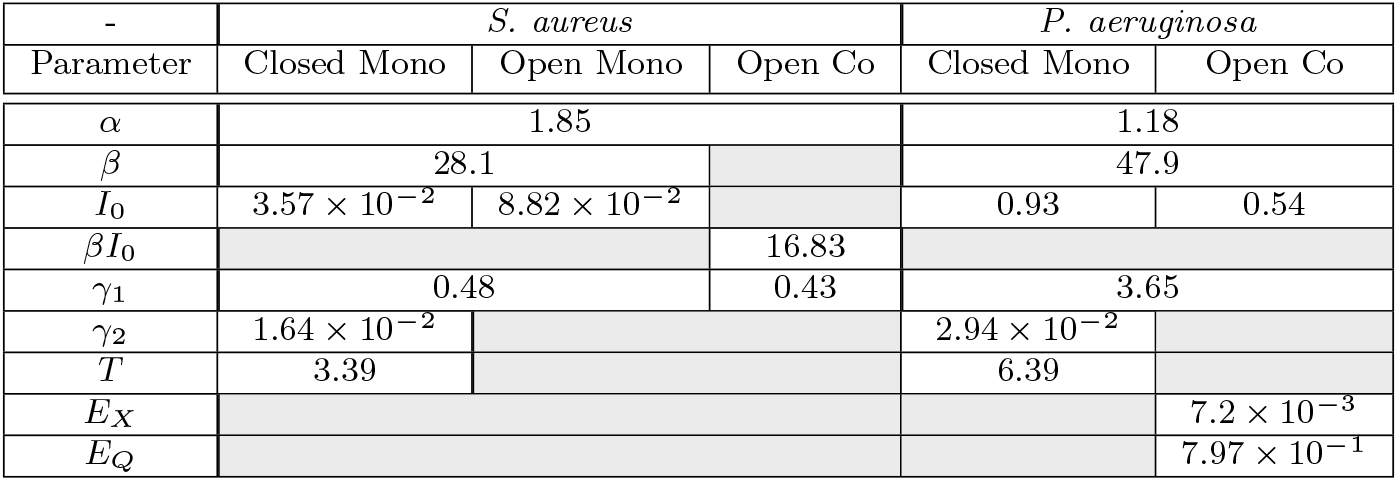
Parameter fitting of Eq. (4) and Eq. (10) using least-squares regression, for *S. aureus* grown a closed environment in monoculture, and in an open environment after being grown in monoculture or coculture with *P. aeruginosa*, and for *P. aeruginosa* grown a closed environment in monoculture, and in an open environment after being grown in coculture with *S. aureus*.

The utility of a model must be substantiated by comparison with previously established models: In service to this, we compare the fitting of the geometric model to fittings of logistic, Gompertz and kinetic models (Fig. 3). The logistic and Gompertz models are presented in Eq. (13).

**Fig. 3:**
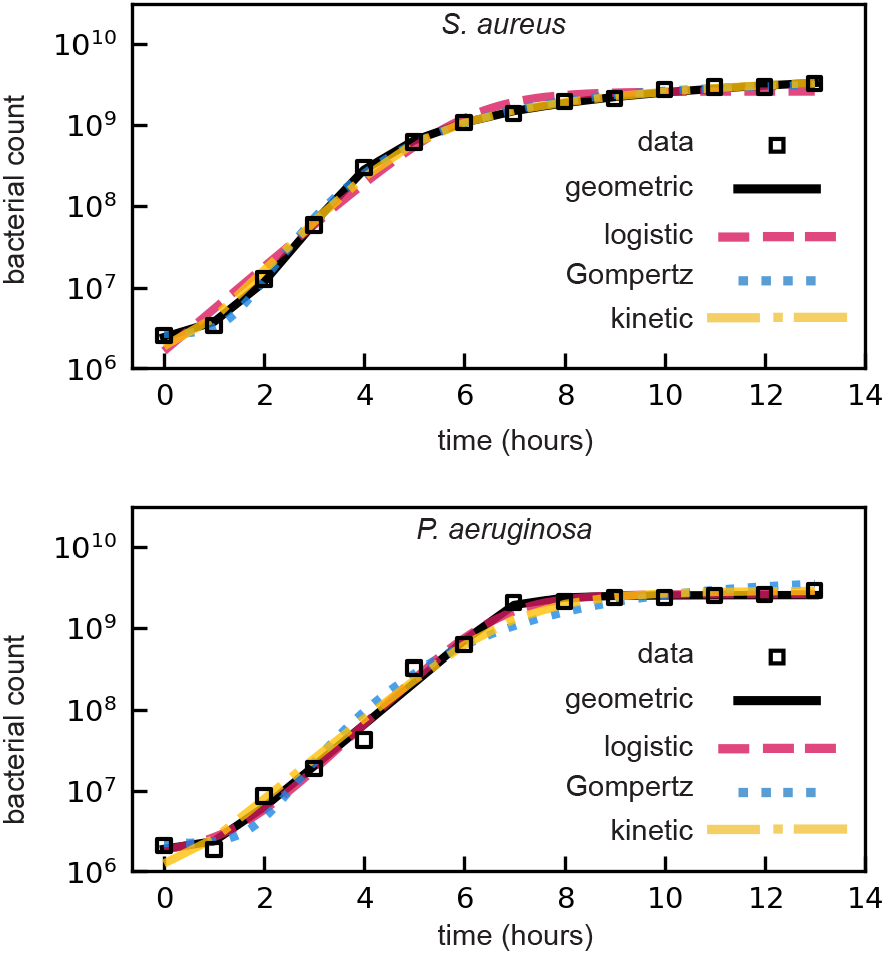
Comparisons of the fits of the Geometric model (solid lines), the Logistic function (dashed), the Gompertz function (dashed), and a kinetic-sense model (dashed-dotted) for both *S. aureus* (upper) and *P. aeruginosa* (lower). The least-square cost for each set of parameter estimations is in Table 2.

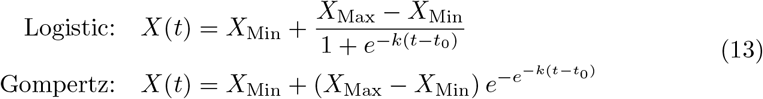

Each of these simple functions requires four parameters which are fitted via least-squares regression. *X*_Min_ and *X*_Max_ can be deduced from the data, *t*_0_ corresponds to the time of maximum slope in growth, and *k* is a measure of the maximum slope (see Appendix C). Parameter estimation results indicate that the geometric model proposed in this study exhibits superior accuracy (as evidenced by a lower cost value, see Table. 2). Furthermore, we recognise that the duration of the lag phases of the logistic and Gompertz functions are influenced by *X*_Max_, even though bacteria ought to be unaware of their carrying capacity until nutrients become limiting: this observation exemplifies the limitations of employing these empirical functions in scenarios where the carrying capacity has not been reached, such as in the open system.

**Table 2:** Cost function of each fit demonstrated in Fig. 3 for both *S. aureus* and *P. aeruginosa*.

| Fit | Least-squares Cost Value |  |
| --- | --- | --- |
|  | *S. aureus* | *P. aeruginosa* |
| Geometric | $4.56 \times 10^{-3}$ | $5.52 \times 10^{-2}$ |
| Logistic | $7.45 \times 10^{-2}$ | $6.65 \times 10^{-2}$ |
| Gompertz | $9.67 \times 10^{-3}$ | $1.73 \times 10^{-1}$ |
| Kinetic | $2.95 \times 10^{-2}$ | $1.14 \times 10^{-1}$ |

Existing kinetic-based models are also commonly employed to describe bacterial growth, treating cells as autocatalysts that interact with external substrates (e.g. glucose, of concentration *S*_*e*_(*t*)) to form a cell–substrate complex of concentration *C*(*t*), which subsequently generates additional biomass (for example, Ref. [30]). The corresponding kinetic equation is:

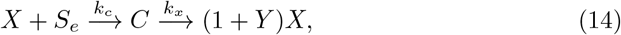

where *k*_*c*_ is the rate of complex formation (analogous to ingestion), *k*_*x*_ is the rate of biomass formation (analogous to assimilation), and *Y* is the number of new cells produced per *μmol* of glucose ingested [30]. The corresponding system of differential equations is given by:

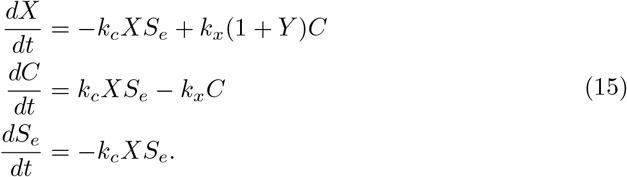

Following parameter estimation via least squares (see Appendix D for the methodology for initial guess estimation), we obtain the fit shown in Fig. 3 (right).

To evaluate the quality of the fits, we compare the cost function obtained from the least-squares algorithm: notably, the geometric model consistently yields the lowest cost across all cases, with a cost value ranging from factors 0.83 to 0.061 times smaller than the next most accurate model.

## 4 Discussion And Conclusion

This paper presents a geometrical model of bacterial growth, grounded in a mechanistic framework that views the cell as a system of interdependent, cascading sub-processes. The model effectively describes the different phases of growth observed in bacterial growth dynamics [1–3]. A particular strength of our model is the description of the lag phase via mechanistic, yet general processes. Furthermore, the model parameters can easily be directly inferred from the data, leading to improved fitting accuracy over existing models.

Naturally, more involved mechanistic models which account for known cellular properties and processes will yield more insightful predictions (for example, Refs. [23–26]). The purpose of this model is to enable comparison: it offers a simple, mechanistic formulation with parameters that can be intuitively interpreted and compared across experiments, shedding light on the cellular stress history (via the parameter *I*_0_). For example, robust model–experiment comparisons could be achieved by more precise experimental estimation of *β* and *γ*_1_ for individual bacterial strains for specific stresses such as the oxidative stress, nutrient limitation, toxin build-up. Resumption of growth in an optimal environment will yield a growth curve from which *β* and *γ*_1_ can be inferred, thus allowing us to infer the source of cellular stress: this could have application in the diagnostic context.

This work has predominantly focussed on capturing the first three phases of bacterial growth: thus, a natural extension of our work is to go on to account for the death phase. The bacterial death phase can be due to a variety of factors, from oxidative stress [39, 40], to altruistic apoptosis [41], to toxicity due to antagonism with other species such as the elastase LasA secreted by *P. aeruginosa* [42] which causes lysis of *S. aureus* [36]. The geometrical model presented in this paper emphasises bacterial growth as a function of components within the cell: oxidative stress and apoptosis could be mapped in this context, as these are consequences of the internal state of the cell. On the other hand, in the context of external factors prompting death, there may need to be phenomenologically unique methods of capturing those interactions: in Ref. [10] a minimal coupling was introduced to describe how chemicals secreted by *P. aeruginosa* prompt the lysis of *S*.*aureus*, for example. Furthermore, the models described in this paper could be further extended to describe bacterial growth in the presence of antibiotics [43, 44] or phage [45, 46] or to describe cellular senescence [47, 48] both in closed and open systems.

More exotic features of cellular growth could be accounted for within this modelling framework by investigating alternatives forms of *I*(*t*), the proportion of inactive components within the cell. One motivating example could be to consider how diauxic growth could be described: there is much research into modelling how and why different metabolites are sequentially metabolised [49–51], and this could be easily incorporated into a consideration of how switching metabolites affects the rate of component inactivation.

## 5 Acknowledgments

## Author contributions

TT: conceptualisation, model formalisation, formal analysis, software writing and maintenance, writing - first draft. UL: data curation, writing - review and editing. SP: data curation, writing - review and editing, supervision. KTA: formal analysis, writing - review and editing, supervision.

## Data and materials availability

All code used to reproduce the analysis presented in this paper can be accessed at Ref. [52].

## Appendix A Corrections in the well-mixed regime

In formulating the geometric model and constructing the interaction space, it is implicitly assumed that if a component shares a downstream process with another, it is likely to share many downstream processes with that component. In other words, the geometric model assumes spatio-temporal correlations between components, and this results in the rate of growth tending to *e*^*-β*^ instead of 0 as the proportion of inactive components tend to 1.

Let us instead assume that in the case of all components being active, a given downstream reaction is dependent upon a random number of components, at least one, with probability *p*. If any of the components are rendered inactive, then that particular process doesn’t happen.

This is directly analogous to a system of *n* switches (components) connected to *N* lights (downstream processes), with each light wired to each switch with probability *p*, and the requirement that each light is connected to at least one switch. The equivalent question is now: given a proportion *I* of switches are flipped off, what is the proportion of lights still turns on (i.e. what is the probability that a given light is still turned on)?

Consider a single light: it is connected to each switch with probability *p*, but must be connected to at least one. We must therefore evaluate *P*_*On*_, which is:

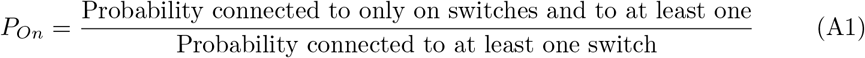

The probability of being connected to no switches is (1 − *p*)^*n*^, thus the probability of being connected to at least one switch is 1 (1 *p*)^*n*^. If *s* are switched off, we need to ensure it is not connected to any of these: the probability of being not connected to a turned-off switch is (1 *-p*)^*s*^, and the probability that it must be connected to one of the remaining *n - s* lights is 1 *-* (1 *-p*)^*n-s*^. Thus:

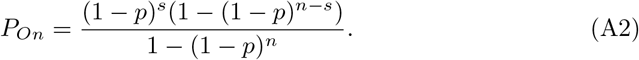

In the limit where there are a large number of switches, we can employ an exponential approximation:

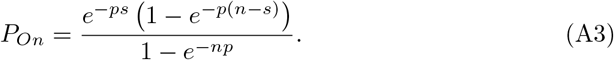

From here we can recognise that *I* = *s/n*, and define the variable λ = *np* as the mean number of connections:

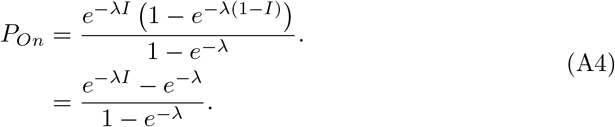

We note that this formulation is an affine transformation of our geometric model, and does tend to 0 as *I* tends to 1.

Thus, the rate equation for bacterial growth would instead resemble:

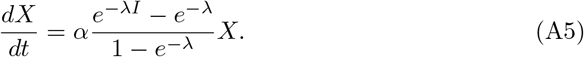

We note that the quantitative features of the lag-exponential transition have identical analytical forms to the original geometric model, thus the values *β* and *γ*_1_ ought to be estimated simultaneously between the models.

## Appendix B Geometric Model Parameter Estimation

For each species, we can estimate the maximum growth rate, *α*, from the wealth of literature regarding the doubling time of each species, *τ*, via the relation *α* = ln(2)*/τ*. Using simply the data, we can obtain an estimate as the maximum value of the slope of the natural log of the data: max 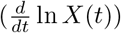.

For the *S. aureus* we fit the closed monoculture, open monoculture and open co-culture data at the same time. We can use a by-eye (or in other words purely data-driven) approximations of the lag time (*t*_lag_), the width of the transition from the lag to the exponential phase (*w*_lag_) the time at which the exponential phase transitions to the stationary phase (*T*), the time at which the stationary phase is approximately achieved (*t*_sat_), and the width of the transition to the stationary phase (*w*_sat_), along with the following relations falling out from our model:

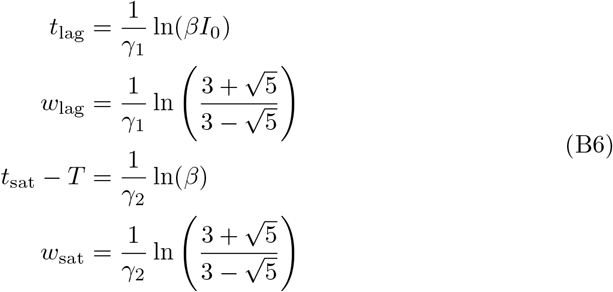

### B.1 *S. aureus* Closed and Open Monoculture

From the closed system, we estimate by eye the parameters of Table B1, and present the derived variables in Table B2.

**Table B1:** Estimates of observed quantities for the case of *S. aureus* in a closed environment.

|  |  |
| --- | --- |
| $t_{\text{lag,closed}}$ | 1 |
| $t_{\text{lag,open}}$ | 0.1 |
| $w_{\text{lag,closed}}$ | 1 |
| $w_{\text{lag,open}}$ | 1 |
| $T$ | 4 |
| $t_{\text{sat}}$ | 8 |
| $w_{\text{sat}}$ | 8 |

**Table B2:**
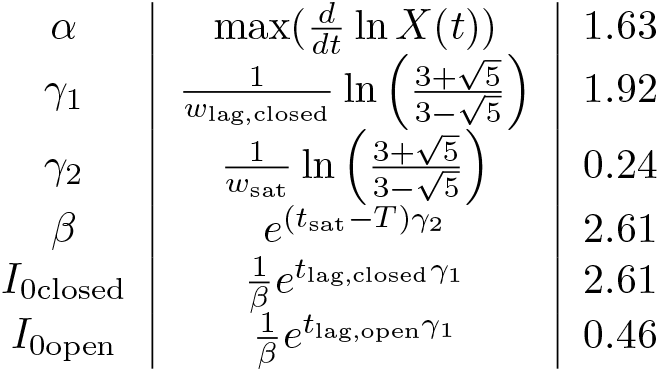
Estimates of derived quantities for the case of *S. aureus* in a closed environment, using the observed quantities in Table B1.

### B.2 *S. aureus* Open Coculture

Similarly, estimate the parameters in Table B3 by eye, from which we derive the parameters in Table B4.

**Table B3:** Estimates of observed quantities for the case of *S. aureus* in an open environment.

|  |  |
| --- | --- |
| $t_{\text{lag}}$ | 6 |
| $w_{\text{lag}}$ | 4 |

**Table B4:**
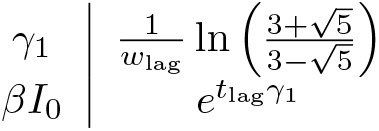
Estimates of derived quantities for the case of *S. aureus* in an open environment, using the observed quantities in Table B3.

### B.3 *P. aeruginosa* Closed and Open

For this system, we also need to find an initial estimate for *E*_*Q*_ and *E*_*X*_. We need to employ by-eye approximation of the period of oscillation, *P*, and the apparent steady state of the open system, *X*^*∗*^, and relate them to the parameters via the relations:

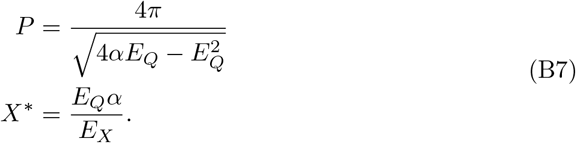

From the closed system, we estimate by eye in Table B5: we note that *t*_lag_ and *w*_lag_ are the same in both examples. The corresponding derived parameters are in Table B6.

**Table B5:** Estimates of observed quantities for the case of *P. aeruginosa* in open and closed environments.

|  |  |
| --- | --- |
| $t_{\text{lag}}$ | 1 |
| $w_{\text{lag}}$ | 1 |
| $T$ | 7 |
| $t_{\text{sat}}$ | 8 |
| $w_{\text{sat}}$ | 1 |
| $P$ | 8 |
| $X^*$ | 100. |

## Appendix C Logistic and Gompertz Empirical Models: Parameter Estimation

The logistic and Gompertz functions are commonly used to fit bacterial growth curves:

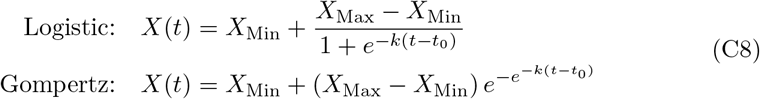

**Table B6:** Estimates of derived quantities for the case of *P. aeruginosa* in open and closed environment, using the observed quantities in Table B5.

|  |  |  |
| --- | --- | --- |
| $\alpha$ | $\max\left(\frac{d}{dt} \ln X(t)\right)$ | 2.03 |
| $\gamma_1$ | $\frac{1}{w_{\text{lag}}} \ln \left( \frac{3+\sqrt{5}}{3-\sqrt{5}} \right)$ | 1.92 |
| $\gamma_2$ | $\frac{1}{w_{\text{sat}}} \ln \left( \frac{3+\sqrt{5}}{3-\sqrt{5}} \right)$ | 1.92 |
| $\beta$ | $e^{(t_{\text{sat}}-T)\gamma_2}$ | 6.85 |
| $I_{0\text{closed}}$ | $\frac{1}{\beta} e^{t_{\text{lag}}\gamma_1}$ | 1.0 |
| $I_{0\text{open}}$ | $\frac{1}{\beta} e^{t_{\text{lag}}\gamma_1}$ | 1.0 |
| $E_Q$ | $\frac{4\alpha \pm \sqrt{16\alpha^2 - \frac{64\pi^2}{P^2}}}{2}$ | 0.317 |
| $E_X$ | $\frac{E_Q \alpha}{X^*}$ | $6.42 \times 10^{-3}$ |

As discussed in the paper, *X*_Min_ and *X*_Max_ are inferred from the data, and *t*_0_ can be inferred by the point of highest slope in each case. *k* can then be deduced from each of the functions and these parameters in Eq. (C9).

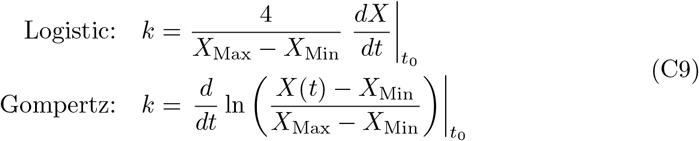

The estimations and corresponding fits for the logistic and Gompertz fits are outlined in Table C7 and Table C8, respectively.

**Table C7:** Comparison of the estimated parameters in the Logistic function Eq. (13), and the fitted parameters obtained by least-squares regression.

| Parameter | *S. aureus* |  | *P. aeruginosa* |  |
| --- | --- | --- | --- | --- |
|  | Estimated | Fitted | Estimated | Fitted |
| $X_{\text{Min}}$ | $2.55 \times 10^6$ | $1.95 \times 10^{-4}$ | $2.12 \times 10^6$ | $1.61 \times 10^6$ |
| $X_{\text{Max}}$ | $2.58 \times 10^9$ | $3.32 \times 10^9$ | $2.93 \times 10^9$ | $2.61 \times 10^9$ |
| $t_0$ | 8 | 6.12 | 6 | 6.67 |
| $k$ | 0.462 | 1.20 | 1.20 | 1.38 |

**Table C8:** Comparison of the estimated parameters in the Gompertz function Eq. (13), and the fitted parameters obtained by least-squares regression.

| Parameter | *S. aureus* |  | *P. aeruginosa* |  |
| --- | --- | --- | --- | --- |
|  | Estimated | Fitted | Estimated | Fitted |
| $X_{\text{Min}}$ | $2.55 \times 10^6$ | $2.67 \times 10^6$ | $2.12 \times 10^6$ | $2.24 \times 10^6$ |
| $X_{\text{Max}}$ | $3.25 \times 10^9$ | $3.32 \times 10^9$ | $2.93 \times 10^9$ | $4.22 \times 10^9$ |
| $t_0$ | 8 | 6.36 | 6 | 7.98 |
| $k$ | 0.217 | 0.404 | 0.940 | 0.337 |

## Appendix D Kinetic-Sense Model: Parameter Estimation

We adopt a kinetic-sense model [30], and estimate parameters using biophysical inferences. To account for the levelling off in the population in mono-culture, we presume that there is a single substrate, of frequency *S*_*e*_, in solution which acts as a limiting factor after it has been consumed by either population of bacteria. We note here that this is in itself an assumption, as there are many possible mechanisms by which a population may level off; for example, when nutrient concentration drops or toxin concentration increases, cells may enter a state of dormancy to survive. As it is impossible to distinguish whether this occurs from a reading of just the cell frequency, we take the simplest approach and assume that the cell growth is inhibited by the absence of nutrients as a leading order behaviour.

We assume that the substrate is imported into the cell at a rate *k*_*c*_ to form a cell-substrate ‘complex’, of frequency *C*, to account for the fact that the importation mechanism for the substrate is limited in how many can be imported at a time. This complex then transitions back to cellular mass as a rate *k*_*x*_, where the new biomass produced is *Y*. The simplest set of equations for a single population grown in monoculture is therefore [30]:

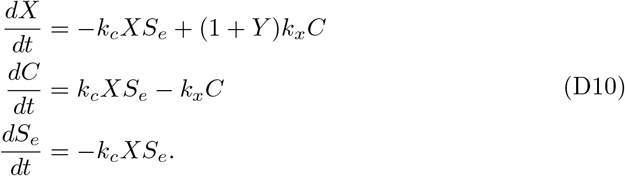

For each population, we can estimate the parameters *k*_*c*_, *Y, K*_*x*_, given the known initial frequencies of cells, *X*_0_ = *X*(0), and the initial frequency of substrate *S*_*e*0_ = *S*_*e*_(0).

*Y* corresponds to the number of cells produced per *μ*mole of the limiting substrate. The simplest assumption is to suggest that the limiting substrate in solution is glucose, of which there is *∼* 0.1*μmol/ml* present [20]. From here, there are two approaches by which we may estimate *Y* for each population.

The theoretical approach is to estimate the direct mass contribution of glucose to the formation of a new cell: this has been discussed that *E. coli* requires 3*×*10^*-*9^*μmol* of glucose as a minimal requirement for replication [53]. We can get an order of magnitude estimation for the amount of glucose required to replicate a *S. aureus* and *P. aeruginosa* by comparison of the relative volumes of these cells with respect to *E. coli*. Given *E. coli* is a rod-shaped bacterium of typical diameter 1*μm* and length 2*μm*, its volume can be approximated as though it were a cylinder, yielding 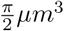. By contrast, *S. aureus* is spherical in nature with an approximate radius of 0.5*μm*, thus a volume 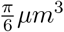, and *P. aeruginosa* is also rod-shaped with a length comparable to *E. coli* yet a diameter of 0.5*μm*, thus an approximate volume of 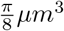. Thus, the volume of each relative to *E. coli* is approximately 3 – 4 times smaller - the order of magnitude of the amount of glucose required to replicate these bacteria would be *∼* 10^*-*9^*μmol*. If it takes 10^*-*9^*μmol* to form one cell, then 1*μmol* would form 10^9^ cells, thus *Y*_*S*_, *Y*_*P*_ *≈*10^9^.

The second approach is to note the conservation law within Eq. (D10):

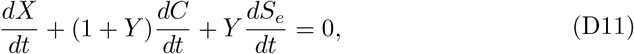

which suggests that *X* +(1 + *Y*)*C* + *Y S*_*e*_ is a conserved quantity. At *t* = 0 we shall assume there has been no nutrient uptake and thus *C*(0) = 0, and *X*(0) = *X*_0_ and *S*_*e*_(0) = *S*_*e*0_ are observed (for *X*_0_) or known (for *S*_*e*0_) quantities. In the limit of *t→ ∞* we see that the populations have levelled this off, and attribute this to the lack of the substrate: Therefore, in this limit lim_*t→∞*_ *C*(*t*) = *S*_*e*_(*t*) = 0, and lim_*t→∞*_ *X*(*t*) = *X*_*∞*_ can be observed. Therefore, by comparing this conserved quantity at *t* = 0 and *t → ∞*:

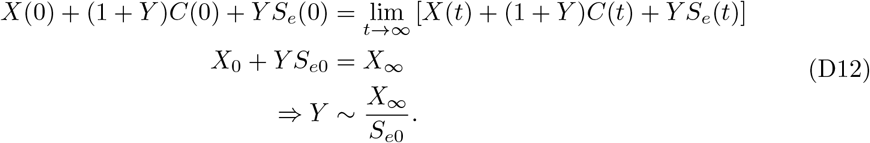

For each population in mono-culture, the long-term frequency of bacteria tend to *∼*10^9^, thus the expectation that *Y ∼*10^10^. Both approaches disagree within only an order of magnitude, thus we will discuss the limiting substrate as though it were glucose. For the same of parameter estimation, we shall employ the conservation law approximation, as it is informed by data specifically.

A mechanistic determination of *k*_*c*_ requires complete knowledge of the rate of glucose ingestion, from the time taken for a single molecule to be imported, and the number of importation proteins: in lieu of this, we can instead assume that the levelling off in population occurs when the rate of glucose consumption is of an order of magnitude similar to the concentration of glucose: *k*_*c*_*XS*_*e*_ *∼ S*_*e*_. Levelling off occurs consistently between populations at *X ∼* 10^9^, thus we anticipate *k*_*c*_ *≈* 10^*-*9^.

*k*_*x*_ can be inferred from the other parameters. First, we consider the evolution of the cell population:

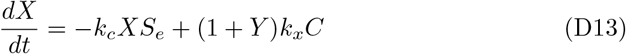

During the exponential phase, the population grows exponentially with some rate constant, *r*. We also assume that during this time, the concentration of substrate is close to the initial concentration: *S*_*e*_ *≈S*_*e*_(0). This means that during the exponential phase,

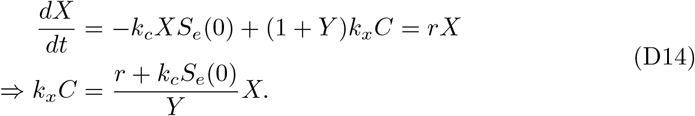

With this identity, we can rearrange the equation for the evolution of the cell-substrate complex:

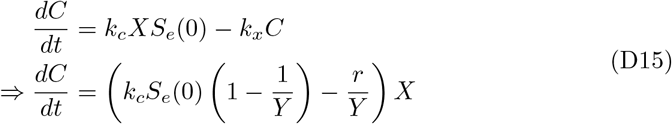

In the exponential phase, we can say that 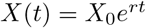, and we assume that during this time *S*_*e*_ *≈* _*e*_(0). Given that the exponential phase spans orders of magnitude, we can choose a timescale to integrate over (*t*_1_, *t*_2_) such that *X*(*t*) = *X*(*t*_2_) *≫ X*(*t*_1_) and *C*(*t*) = *C*(*t*_2_) *≫ C*(*t*_1_) and thus solve

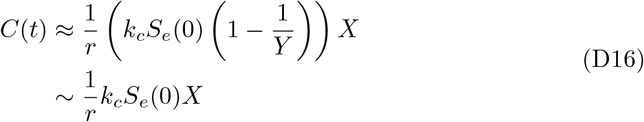

We can substitute Eq. (D14) and Eq. (D16) together, to obtain an expression for *k*_*x*_ in terms of the other parameters:

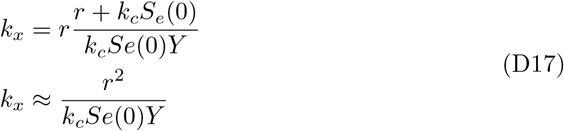

Given *Y ∼* 10^10^, *S*_*e*_(0) *∼* 0.1, *k*_*c*_ *∼* 10^*-*9^ and the observed growth rate is *r ∼* 1, we obtain *k*_*x*_ *∼* 1

**Table D9:** Comparison of the estimated parameters in the kinetic model Eq. (15), and the estimated parameters obtained by least-squares regression.

|  | Estimate | Fitted |  |
| --- | --- | --- | --- |
| Parameter |  | *S. aureus* | *P. aeruginosa* |
| $k_c(\mu\text{mol}/h)$ | $10^{-9}$ | $3.64 \times 10^{-9}$ | $1.02 \times 10^{-9}$ |
| $k_x(h^{-1})$ | 1 | 0.117 | 0.754 |
| $Y$ | $10^{10}$ | $5.08 \times 10^{10}$ | $2.88 \times 10^{10}$ |

